# A highly contiguous genome assembly reveals sources of genomic novelty in the symbiotic fungus *Rhizophagus irregularis*

**DOI:** 10.1101/2022.10.19.511543

**Authors:** Bethan F. Manley, Jaruwatana S. Lotharukpong, Josué Barrera-Redondo, Gokalp Yildirir, Jana Sperschneider, Nicolas Corradi, Uta Paszkowski, Eric A. Miska, Alexandra Dallaire

## Abstract

The root systems of most plant species are aided by the soil foraging capacities of symbiotic Arbuscular Mycorrhizal (AM) fungi of the Glomeromycotina subphylum. Despite recent advances in our knowledge of the ecology and molecular biology of this mutualistic symbiosis, our understanding of the AM fungi genome biology is just emerging. Presented here are the most contiguous and highest-quality nuclear and mitochondrial genome assemblies of an arbuscular mycorrhizal fungus to date, achieved through Nanopore long-read DNA sequencing and Hi-C data. This haploid genome assembly of *Rhizophagus irregularis*, alongside short- and long-read RNA-Sequencing data, was used to produce a comprehensive annotation catalogue of gene models, repetitive elements, small RNA loci, and DNA cytosine methylome. A phylostratigraphic gene age inference framework revealed that the birth of genes associated with nutrient transporter activity and transmembrane ion transport systems predates the emergence of Glomeromycotina. While symbiotic nutrient cycling in AM fungi relies on genes that existed in ancestor lineages, a burst of Glomeromycotina-restricted genetic innovation is also detected. Analysis of the chromosomal distribution of genetic and epigenetic features highlights evolutionarily young genomic regions that produce abundant small RNAs, suggesting active RNA-based monitoring of genetic sequences surrounding recently evolved genes. This chromosome-scale view of the genome of an AM fungus genome reveals previously unexplored sources of genomic novelty in an organism evolving under an obligate symbiotic life cycle.

**Highlights:**

- Assembly of 32 highly contiguous chromosomal scaffolds for *R. irregularis*, with 23 complete and gapless
- Gene annotation based on short- and long-read RNA-Seq data from different developmental stages
- Complete annotation set including mitochondrial genes, DNA methylome, small RNAome, repetitive/transposable elements, functional annotation
- Identification of a burst of lineage-restricted genetic innovation in the Glomeromycotina subphylum

## Introduction

Uprooting almost any terrestrial plant reveals the arbuscular mycorrhizal (AM) symbiosis, a mutually beneficial interaction between most land plant species and members of the fungal Glomeromycotina subphylum (Parniske, 2008). AM fungi are multinucleate, obligate symbionts that exist in all terrestrial ecosystems (Davison et al., 2015) and engage in symbioses with a wide range of plant species, often simultaneously (Bever, 2002). While ecological and molecular mechanistic evidence suggest that the AM symbiosis relies on reciprocal transfer of organic and inorganic nutrients through a permeable membranous interface (Bonfante & Genre, 2010), our understanding of the genomic basis of this symbiotic lifestyle remains limited by the fact that whole-genome sequencing data is available for a limited number of AM species (Kobayashi et al., 2018; Malar et al., 2021; Montoliu-Nerin et al., 2021; Morin et al., 2019; Prasad Singh et al., 2019; Sahraei et al., 2022; Singh et al., 2021; Sun et al., 2019; Trepanier et al., 2005; Venice et al., 2020). These include genome assemblies of multiple isolates of the model species, *Rhizophagus irregularis*, and the homokaryotic laboratory strain DAOM197198 (**Figure 1**)(Chen, Mathieu, et al., 2018; Chen, Morin, et al., 2018; Lin et al., 2014; Maeda et al., 2018; Tisserant et al., 2013; Yildirir et al., 2022). The most recent genome assembly of DAOM197198 represented a sizeable step-up in genome contiguity and quality (Yildirir et al., 2022), however, the contig N50 of 2.3Mb and quantity of gaps in this assembly is lagging behind recent fungal genome assemblies (Chung et al., 2021; Liu et al., 2020). The *de novo* assembly of a reference genome is a crucial step for the genetic research of a given organism. To best support genomic and transcriptomic research, the ideal resource is a fully sequenced, contiguous genomic assembly with few gaps (Church et al., 2011; Rhie et al., 2021). Recent attention has been paid to the contribution of epigenetic and transposable element landscapes of *R. irregularis* to the adaptation and evolution of this species (Chaturvedi et al., 2021; Dallaire et al., 2021; Yildirir et al., 2022). The production of a higher quality genome for *R. irregularis* will further enable research into the repetitive landscape and the genomic organisation of Glomeromycotina fungi and their relatives, providing crucial insights into the biology and evolutionary history of the AM symbiosis.

**Figure 1.**
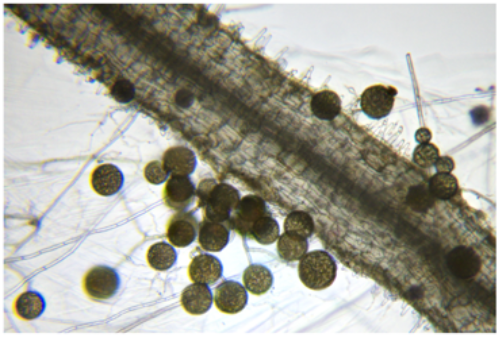
Carrot root with extraradical hyphae and spores of *R. irregularis*.

This study presents a highly contiguous and near-gapless long-read assembly of the *R. irregularis* isolate DAOM197198 achieved using long Nanopore reads, Hi-C data, and manual curation. Nanopore RNA-Sequencing was generated for *R. irregularis*, producing long reads that span entire transcripts to guide and improve Illumina short-read-based gene model predictions, and to enable the annotation of untranslated (UTR) regions, prediction of poly(A) signal and poly(A) tail length analyses. Repetitive element, small RNA loci and DNA cytosine methylome annotations are also provided. These datasets were combined with a tree-of-life scale analysis of gene birth events (Barrera-Redondo et al., 2022), which assigns an evolutionary age to protein-coding genes of *R. irregularis* and identifies taxonomically-restricted genes that have no detectable homologs in other organisms. This analysis identifies molecular functions that are ancestral to the Glomeromycotina and describes an important gene birth event coinciding with their emergence. The chromosomal distribution of genetic and epigenetic features uncovers evolutionarily young regions of the genome that are potential cradles for new genes and small RNA production.

## Results

### De novo assembly of the *R. irregularis* genome

Assembly using trimmed Nanopore reads resulted in 44 contigs that were polished using Illumina reads (Maeda et al. 2018). Two of these contigs were filtered out due to their size of <500bp, resulting in a polished and filtered assembly of 42 contigs (**Figure S1A**). The assembly process produced a complete, circular mitochondrial genome of 70,793 bp, within the size range of other AM fungal mitochondrial genomes (**Figure 2A**) (Lee & Young, 2009; Nadimi et al., 2016). This mitochondrial genome was annotated using MitoHifi and contains sequences encoding transfer RNAs (tRNAS), ribosomal subunits and genes typically identified on a fungal mitochondrial genome. Manual curation based on Hi-C read alignment to the nuclear genome assembly was used to assign the remaining 42 contigs to 32 chromosomal units (**Figure 2B**). Prior to manual curation, contig N50 and L50 were 3,900,757 bp (~3.9 Mb) and 15, respectively, rising to a scaffold N50 and L50 of 5,085,394 (~5 Mb) and 13 post-curation (**Table 1**). 23 of these scaffolds were complete and gapless chromosomes (**Figure 2C**). 17 of the 32 chromosome-scale scaffolds of *R. irregularis* were produced telomere-to-telomere, with telomeric repeats of sequence TTAGGG_n_ identified at both 5’ and 3’ ends of the scaffolds, and an additional 14 containing one telomere (**Figure 2C**). Average Illumina and Nanopore read coverage were highly uniform across all scaffolds, indicating that repetitive sequences are fully resolved (**Figure S1B**). The 32 chromosomes display extensive macro-synteny to a recent assembly of this species, except for stretches of chromosomes 1 and 5 (**Figure S1C**). This assembly suggests a misjoin in a previous assembly of this species, which would result in the potential overestimation of the number of *R. irregularis* chromosomes (**Table 1**). Research into the location of centromeric repeats of this symbiotic fungus may aid further analyses into chromosome number and structure of these organisms. The final haploid nuclear assembly following removal of the circular mitochondrial contig is 146,773,001bp in size.

**Figure 2.**
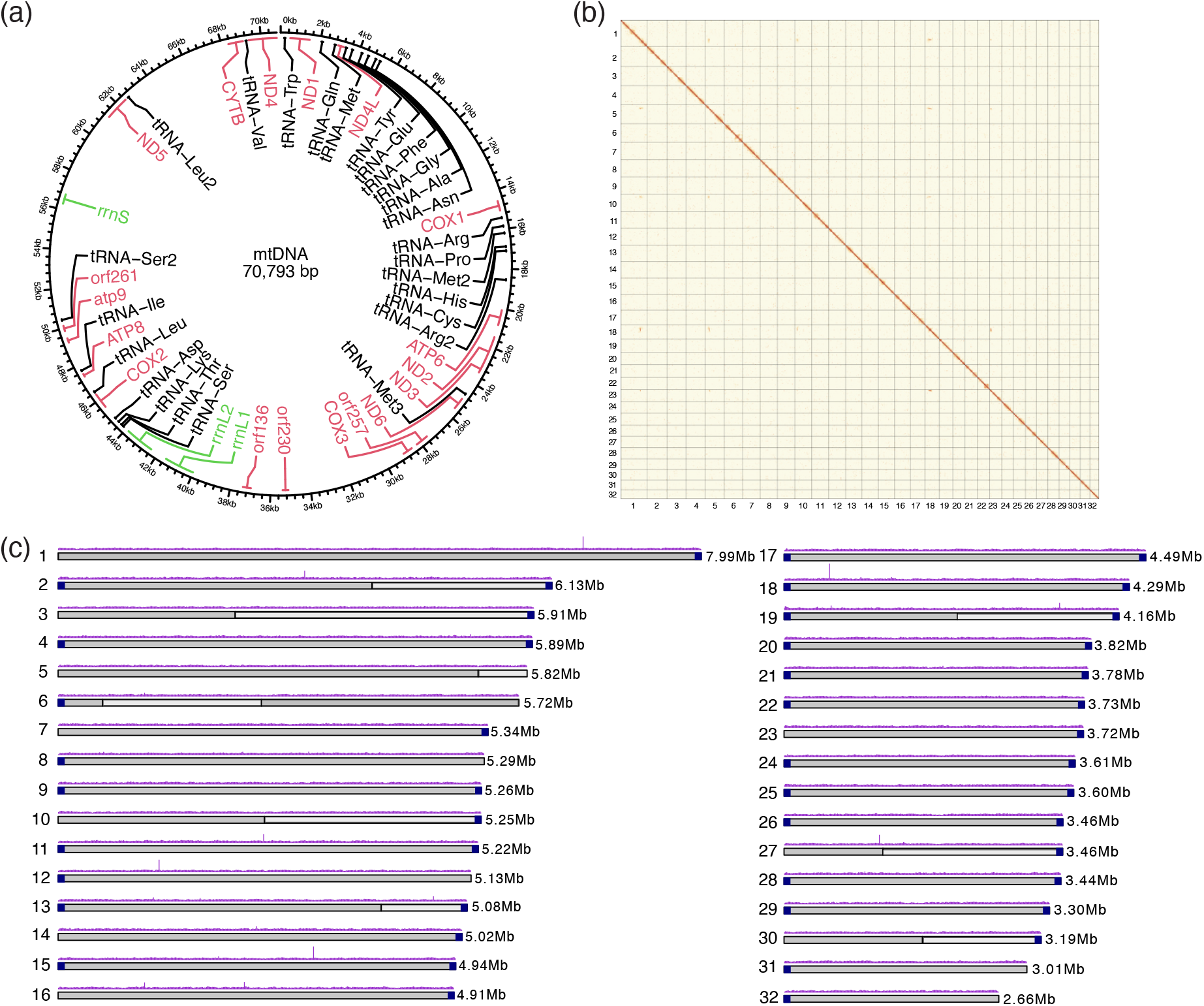
Nuclear and mitochondrial genome assemblies of *R. irregularis*. a) Circular map of mitochondrial genome with annotated genes (pink), tRNAs (black) and rRNAs (green). b) Hi-C contact map visualised in PretextView. Chromosomes are displayed in size order from left to right (1-32). c) Physical map of 32 chromosomes numbered according to size (Mb). Grey colouring of the ideogram highlights contigs that were scaffolded together. Telomeric sequences are represented by dark blue squares at the ends of ideograms. Nanopore read coverage is shown as a purple histogram.

**Table 1.**
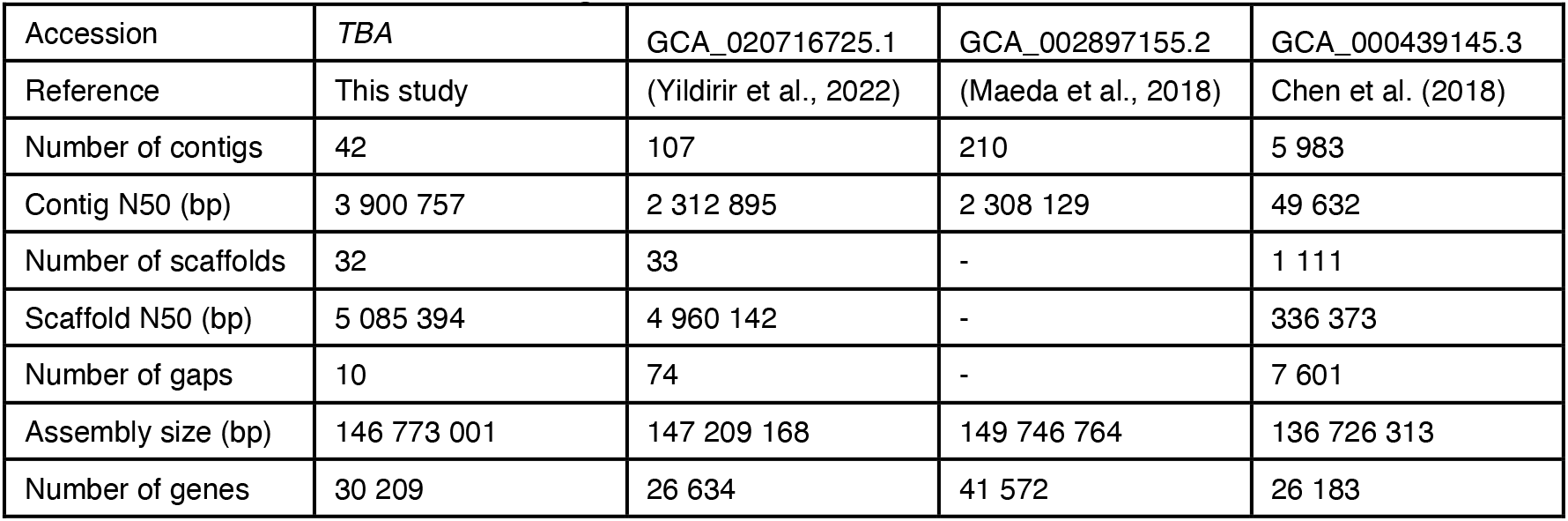
Summary of *R. irregularis* genome assemblies

### Genome annotation using short- and long-read sequencing

Following modelling, curation and masking of repetitive sequences and transposable elements, proteincoding genes were annotated using published Illumina RNA-Sequencing (RNA-Seq) reads from multiple life stages (**Table 2**). This Illumina-based gene annotation was manually curated to remove transposable elements, leaving 30,230 gene models (Illumina-based gene annotation). Gene models were then refined with long Nanopore RNA-Seq reads, improving the support of exon-intron boundaries by sequencing reads (**Figure 3A**; Illumina+Nanopore-based annotation), and increasing gene and exon length (**Figure 3B-C**). Updated gene models were manually curated to remove transposable elements, resulting in a final annotation of 30,209 genes. This gene count is consistent with previous studies into genes encoded by AM fungal genomes (Miyauchi et al., 2020). Long-read data did not change the overall BUSCO score (96.9%) but moved one duplicated BUSCO gene to the single-copy category (**Figure 3D**). Functional annotation of gene models indicated that long reads improved the proportion of genes with assigned Gene Ontology (GO) terms, PFAM domains, InterPro domains and secretion signals, while the number of biosynthetic genes and CAZymes remained constant (**Figure 3E**). Refining gene models with long reads therefore resulted in more accurate gene models and a higher number of functionally annotated genes. Examples of updated gene models include glucosamine-6-phosphate isomerase (NAG1) and Crinkler effector 10 (CRN10), two genes thought to be involved in arbuscule development and function (Kobae et al., 2015; Voss et al., 2018). Compared to previous accessions, long-read data revealed two novel transcript isoforms of NAG1 that contain an additional exon (**Figure 3F**; g17052-T1 and g17052-T2). A misannotated first intron of CRN10 was fixed, and the updated gene sequence is identical to the one described in (Voss et al., 2018).

**Table 2.**
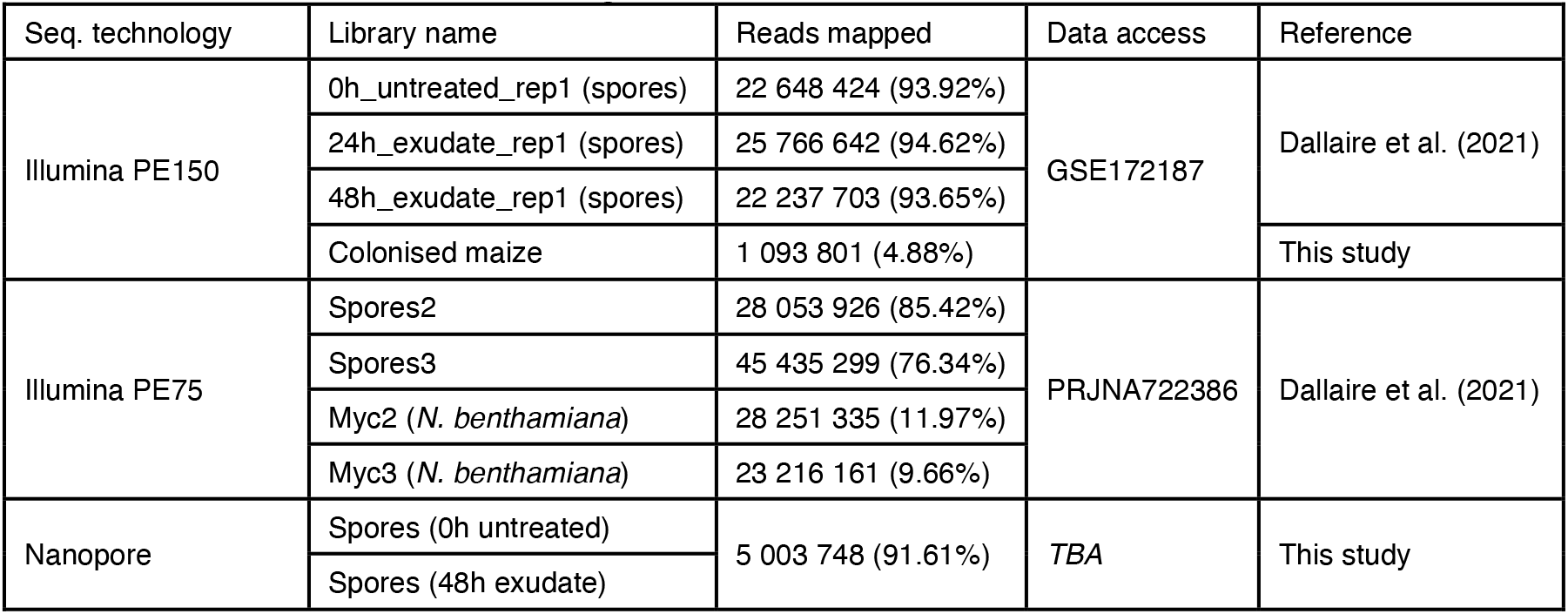
RNA-Seq datasets used for gene annotation

| Seq. technology | Library name | Reads mapped | Data access | Reference |
| --- | --- | --- | --- | --- |
| Illumina PE150 | 0h_untreated_rep1 (spores) | 22 648 424 (93.92%) | GSE172187 | Dallaire et al. (2021) |
|  | 24h_exudate_rep1 (spores) | 25 766 642 (94.62%) |  |  |
|  | 48h_exudate_rep1 (spores) | 22 237 703 (93.65%) |  |  |
|  | Colonised maize | 1 093 801 (4.88%) |  | This study |
| Illumina PE75 | Spores2 | 28 053 926 (85.42%) | PRJNA722386 | Dallaire et al. (2021) |
|  | Spores3 | 45 435 299 (76.34%) |  |  |
|  | Myc2 (*N. benthamiana*) | 28 251 335 (11.97%) |  |  |
|  | Myc3 (*N. benthamiana*) | 23 216 161 (9.66%) |  |  |
| Nanopore | Spores (0h untreated) | 5 003 748 (91.61%) | *TBA* | This study |
|  | Spores (48h exudate) |  |  |  |

**Figure 3.**
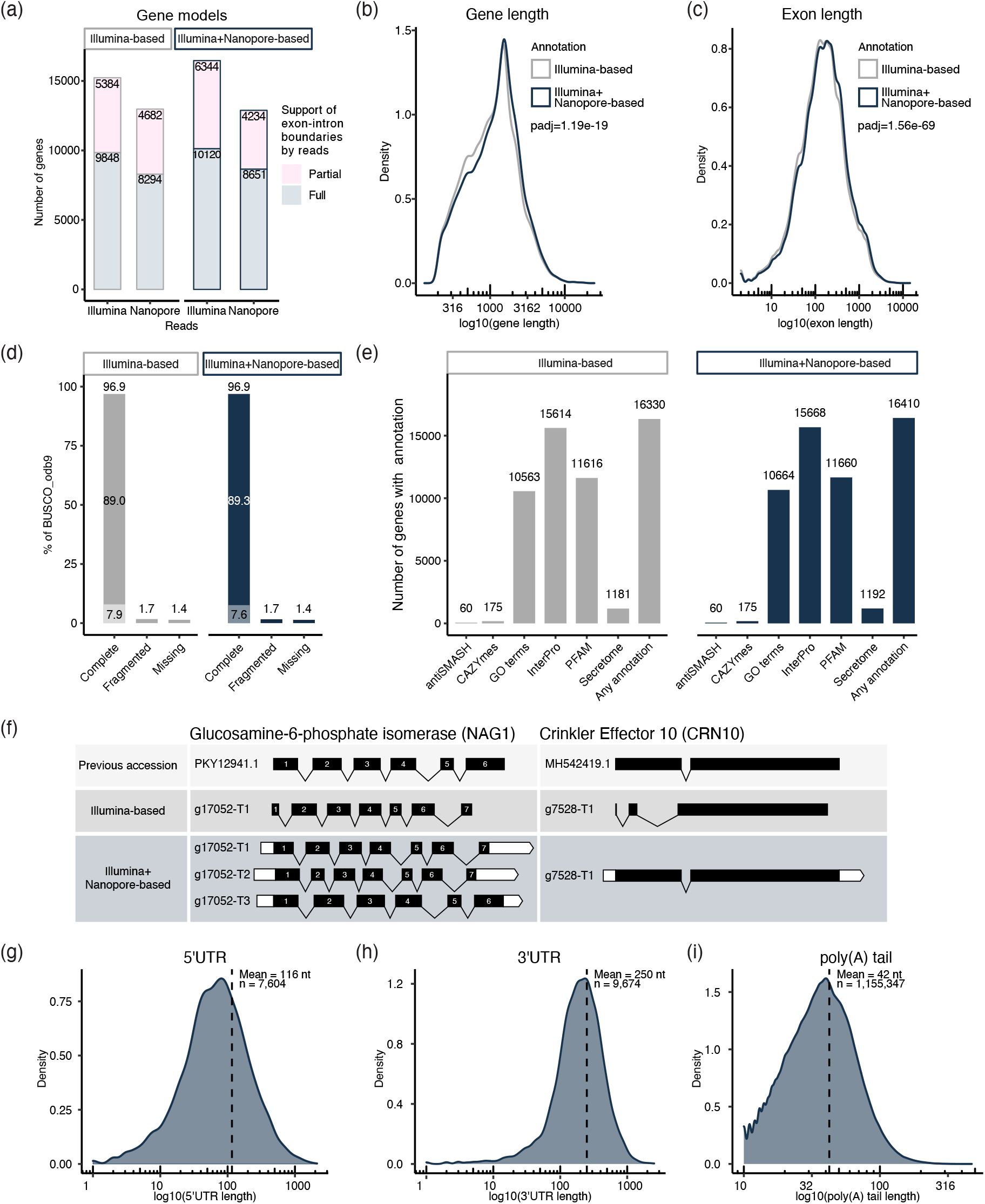
General features of revised gene models. a) Support of exon-intron boundaries of Illumina-based and Illumina+Nanopore-based gene annotations by Illumina and Nanopore RNA-Seq reads. The number of genes with all boundaries (full support) and partial boundaries (partial support) supported by experimental evidence is indicated. Exon-less genes are not displayed. Comparison of gene b) and exon c) length distribution between Illumina and Illumina+Nanopore-based gene annotations. The x axes are on a log scale and a paired T-test was used to assess statistical significance. d) Comparison of BUSCO gene categories. The complete stack is split into single copy (top) and duplicated (bottom). e) Comparison of functional annotation of Illumina-based and Illumina+Nanopore-based gene models. f) Comparison of gene models revised using long-read data to previous annotations and Illumina-based gene models. Black boxes represent exons, lines are introns and white boxes are UTRs. Length distribution of 5UTRs g) and 3’UTRs h) of the Illumina+Nanopore gene models. i) Length distribution of poly(A) tails detected in spores. The x axes are on a log scale.

### Untranslated regions, poly(A) tails and the poly(A) signal of *R. irregularis*

Long RNA-Seq reads provided evidence for untranslated region (UTR) prediction, polyadenylation site detection and poly(A) tail length analyses. 5’UTR and 3’UTR length distributions have respective means of 116 and 250 nucleotides (nt) (**Figure 3G-H**), which are comparable to the fungal averages (134 and 237nt respectively) and within the known ranges of eukaryotic UTR lengths (100-200nt 5’UTR; 200-1000nt 3’UTR) (Bruno et al., 2010; Lin & Li, 2012; Mignone et al., 2002; Pesole et al., 2001). A MEME motif search in the 50bp preceding the poly(A) tails of 242,742 unique poly(A) sites yielded one significantly enriched hexanucleotide motif, the canonical AAUAAA (**Table 3**; E-value 1.3e-24) (Bailey & Elkan, 1994). This sequence accounts for 56.7% of detected poly(A) sites, indicating high sequence conservation to the mammalian polyadenylation signal, compared to yeast (13.2%), *Aspergillus oryzae* (6%), *Arabidopsis thaliana* (10%) and *Oryza sativa* (7%) (Table 3)(Graber et al., 1999; Loke et al., 2005; Shen et al., 2008; Tanaka et al., 2011). Additional derivatives such as AUUAAA and AAUAUA were also detected but were not significantly enriched. The distribution of poly(A) tail lengths in spore transcripts ranged from 10 to 473nt, with a mean of 42nt (**Figure 3I**), which is comparable to the 50nt average observed in *S. cerevisiae* using similar methods (Tudek et al., 2021).

**Table 3.**
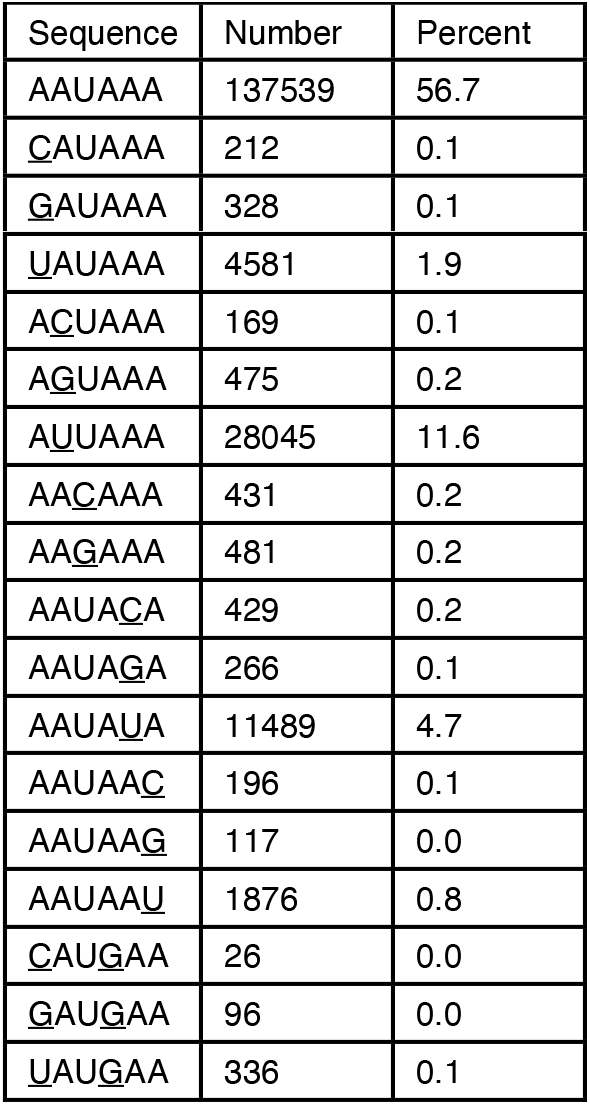
*R. irregularis* polyadenylation signal(s)alic> polyadenylation signal(s)

### A burst of gene novelty with the emergence of Glomeromycotina fungi

A tree of life scale comparative genomics analysis was used to estimate the evolutionary ages of *R. irregularis* genes, tracing gene birth events to the last universal common ancestor (Barrera-Redondo et al., 2022). This analysis suggests that 34% (n=10,250) of *R. irregularis* genes have homologs across taxonomic levels and date back to the origin of cellular organisms (**Figure 4A**; all genes). This most ancient phylorank (phylorank 1) is enriched for basic cellular functions and primary metabolic processes such as transcription, translation and regulation of cell cycle (**Table S1**), which are expected to be conserved across the tree of life. Notably, 2373 out of 2533 members of *R. irregularis’* expanded kinome are found at phylorank 1, consistent with protein phosphorylation as a fundamental mechanism of cell signalling (**Table S1** and **Table S2**)(Kwon et al., 2019). All phosphate transporters (PT1 to PT7), ammonium transporters (AMT1, AMT2, AMT3) and monosaccharide transporters (MST2, MST3, MST4) are found at phylorank 1 (**Table 4**). As may be expected, this analysis suggests that phosphate, nitrogen and carbohydrate efflux and homeostasis are ancestral molecular functions that emerged long before AM fungi. Our analysis revealed comparable numbers of highly conserved genes in the Glomeromycotina fungi *Gigaspora margarita* (40%, n=11,731) and *Geosiphon pyriformis* (46%; n=6875)(**Figure S2A**, phylorank 1). The Glomeromycotina, Mucoromycotina and Mortierellomycotina species analysed here share similar gene age distributions until the emergence of the Mucoromycota, where each lineage displays their independent historical patterns of gene emergence (**Figure S2A**, phyloranks 1 to 5).

**Figure 4.**
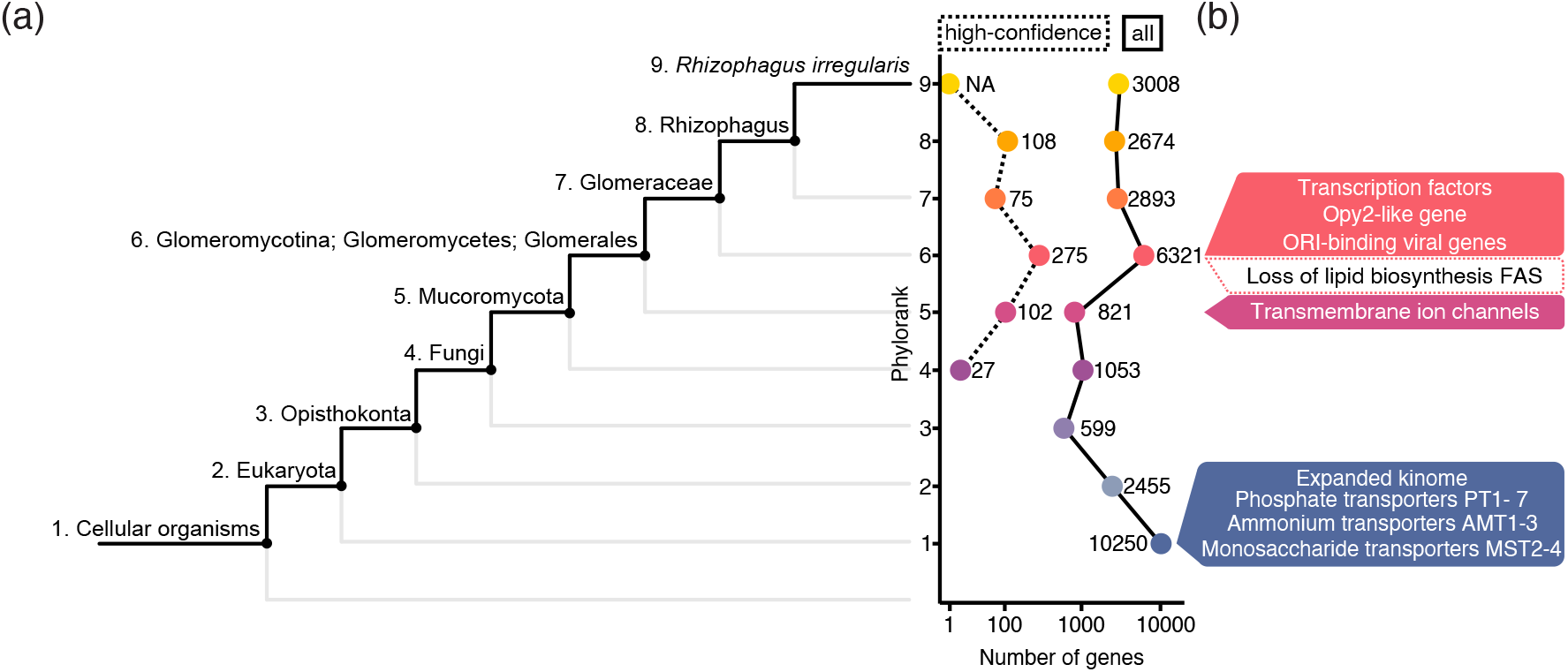
Phylostratigraphy analysis of *R. irregularis* genes. a) Left panel: R. irregularis lineage. Right panel: number of genes at each phylorank before (full line) and after (dashed line) accounting for homology detection failure (HDF). Dashed line represents genes with high confidence phyloranks that could not be explained by HDF. b) Model of gene birth and gene loss in the *R. irregularis* lineage.

**Table 4.** Nutrient transporter genes at phylorank 1

| Function | Gene Name | Gene ID |
| --- | --- | --- |
| Phosphate transporter | PT1 | g11592 |
|  | PT2 | g7615 |
|  | PT3 | g111 |
|  | PT4 | g31083 |
|  | PT5 | g18438 |
|  | PT6 | g27858 |
|  | PT7 | g19437 |
| Ammonium transporter | AMT1 | g16666 |
|  | AMT2 | g1222 |
|  | AMT3 | g18142 |
| Monosaccharide transporter | MST2 | g24501 |
|  | MST3 | g19549 |
|  | MST4 | g26862 |

A gene birth event was detected at the Glomeromycotina phylorank, indicating that their emergence is marked by a burst of lineage-restricted evolutionary novelty (**Figure 4A**; phylorank 6; all genes). One caveat of phylostratigraphy is that gene age is often underestimated because of the inability of pairwise aligners to trace back homologs in outgroups that are too evolutionarily distant. Robust assessment of gene birth events therefore relies on testing the null hypothesis of homology detection failure (HDF) in order to achieve high-confidence predictions (Barrera-Redondo et al., 2022). A more stringent analysis taking into account HDF of recently evolved genes confirmed the burst of gene birth in Glomeromycotina (**Figure 4A-C**; phylorank 6; high confidence). Confidently ranked genes born in the Glomeromycotina include an HTH APSES-type transcription factor (g4815), a Zn(2)-C6 fungal-type transcription factor (g25112), an Opy2-like membrane anchor protein (g2640), two uncharacterised Crinkler-type effectors (g11050, g27662), a Complex 1 LYR protein (g6617), and many F-box and Leucine repeat genes (**Figure 4C**, **Table S2** and **Table S3**). Two GO terms related to replication were enriched at the high confidence phylorank 6, and are linked to genes with a potential viral origin (**Figure 4C**, **Table 5**). These genes have putative replication-origin binding domains (InterPro domain IPR003450), and were most likely acquired through horizontal transfer in the common ancestor of Glomeromycotina and subsequently inherited vertically throughout the whole lineage. Genes born at the emergence of Glomeromycotina may encode functions that were crucial for their evolutionary success and diversification, such as developmental innovation for symbiosis or obligate biotrophy.

**Table 5.**
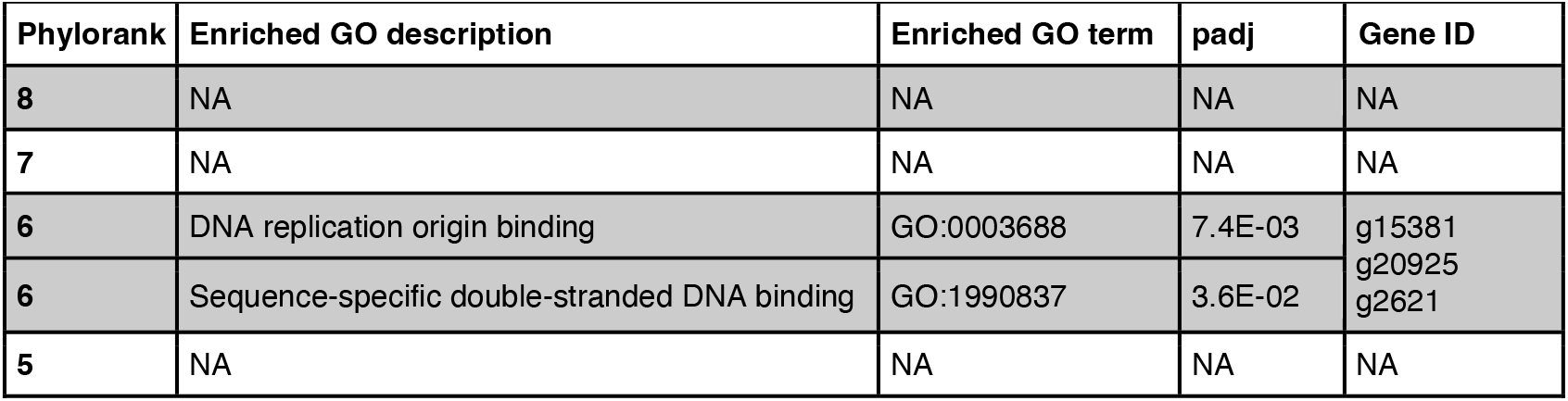
GO terms enriched in genes with high confidence phyloranks

Although the ages of most genes at the Mucoromycota phylorank may be underestimated without accounting for HDF, general shifts in protein sequence space can still be captured (Domazet-Lošo et al., 2022). GO term enrichment analyses were performed to investigate molecular functions that are ancestral to Glomeromycotina. GO terms related to ion transport, transmembrane transporter activity and membrane components are significantly enriched at the Mucoromycota phylorank (**Table S1**; phylorank 5). The genes underlying this enrichment mainly consisted of a group of 32 transient receptor channel subfamily V-like genes with predicted permeability to Ca^2+^ (**Figure 4B**, **Table 6**) (Nilius & Szallasi, 2014). Innovation in transmembrane ion transport therefore precedes Glomeromycotina and may be a feature that marked the evolutionary transition of Mucoromycota fungi.

**Table 6.** Genes with enriched membrane and ion transport GO terms at Mucoromycota phylorank 5

| GO term ID | Description | p.val | FDR | Genes |
| --- | --- | --- | --- | --- |
| GO:0005216 | ion channel activity | 4.4E-35 | 4.4E-35 | g12620, g14470, g14472, g14478, g17590, g17616, g17700, g17949, g22276, g22280, g22289, g22304, g22306, g22308, g22310, g22315, g22331, g22332, g22338, g22346, g22348, g22357, g22367, g22372, g22476, g22485, g22497, g25077, g25094, g31340, g6647, g6926 |
| GO:0022803 | passive transmembrane transporter activity | 1.5E-33 | 1.5E-33 | g12620, g14470, g14472, g14478, g17590, g17616, g17700, g17949, g22276, g22280, g22289, g22304, g22306, g22308, g22310, g22315, g22331, g22332, g22338, g22346, g22348, g22357, g22367, g22372, g22476, g22485, g22497, g25077, g25094, g31340, g6647, g6926 |
| GO:0015267 | channel activity | 1.5E-33 | 1.5E-33 | g12620, g14470, g14472, g14478, g17590, g17616, g17700, g17949, g22276, g22280, g22289, g22304, g22306, g22308, g22310, g22315, g22331, g22332, g22338, g22346, g22348, g22357, g22367, g22372, g22476, g22485, g22497, g25077, g25094, g31340, g6647, g6926 |
| GO:0006811 | ion transport | 8.5E-32 | 8.5E-32 | g12620, g14470, g14472, g14478, g17590, g17616, g17700, g17949, g22276, g22280, g22289, g22304, g22306, g22308, g22310, g22315, g22331, g22332, g22338, g22346, g22348, g22357, g22367, g22372, g22476, g22485, g22497, g25077, g25094, g31340, g6647, g6926 |
| GO:0015318 | inorganic molecular entity transmembrane transporter activity | 1.4E-28 | 1.4E-28 | g12620, g14470, g14472, g14478, g17590, g17616, g17700, g17949, g22276, g22280, g22289, g22304, g22306, g22308, g22310, g22315, g22331, g22332, g22338, g22346, g22348, g22357, g22367, g22372, g22476, g22485, g22497, g25077, g25094, g31340, g6647, g6926 |
| GO:0015075 | ion transmembrane transporter activity | 1.4E-26 | 1.4E-26 | g12620, g14470, g14472, g14478, g17590, g17616, g17700, g17949, g22276, g22280, g22289, g22304, g22306, g22308, g22310, g22315, g22331, g22332, g22338, g22346, g22348, g22357, g22367, g22372, g22476, g22485, g22497, g25077, g25094, g31340, g6647, g6926 |
| GO:0031224 | intrinsic component of membrane | 3.9E-22 | 3.9E-22 | g12620, g14470, g14472, g14478, g17590, g17616, g17700, g17949, g22276, g22280, g22289, g22304, g22306, g22308, g22310, g22315, g22331, g22332, g22338, g22346, g22348, g22357, g22367, g22372, g22476, g22485, g22497, g25077, g25094, g28535, g31340, g6647, g6778, g6926, g7404, g7406, g7408, g9272, g9509 |
| GO:0016021 | integral component of membrane | 3.9E-22 | 3.9E-22 | g12620, g14470, g14472, g14478, g17590, g17616, g17700, g17949, g22276, g22280, g22289, g22304, g22306, g22308, g22310, g22315, g22331, g22332, g22338, g22346, g22348, g22357, g22367, g22372, g22476, g22485, g22497, g25077, g25094, g28535, g31340, g6647, g6778, g6926, g7404, g7406, g7408, g9272, g9509 |
| GO:0022857 | transmembrane transporter activity | 9.8E-20 | 9.8E-20 | g12620, g14470, g14472, g14478, g17590, g17616, g17700, g17949, g22276, g22280, g22289, g22304, g22306, g22308, g22310, g22315, g22331, g22332, g22338, g22346, g22348, g22357, g22367, g22372, g22476, g22485, g22497, g25077, g25094, g31340, g6647, g6926 |
| GO:0005215 | transporter activity | 6.6E-19 | 6.6E-19 | g12620, g14470, g14472, g14478, g17590, g17616, g17700, g17949, g22276, g22280, g22289, g22304, g22306, g22308, g22310, g22315, g22331, g22332, g22338, g22346, g22348, g22357, g22367, g22372, g22476, g22485, g22497, g25077, g25094, g31340, g6647, g6926 |
| GO:0016020 | membrane | 6.6E-14 | 6.6E-14 | g12620, g14470, g14472, g14478, g17590, g17616, g17700, g17949, g22276, g22280, g22289, g22304, g22306, g22308, g22310, g22315, g22331, g22332, g22338, g22346, g22348, g22357, g22367, g22372, g22476, g22485, g22497, g25077, g25094, g28535, g31340, g6647, g6778, g6926, g7404, g7406, g7408, g9272, g9509 |
| GO:0006810 | transport | 9.6E-14 | 9.6E-14 | g12620, g14470, g14472, g14478, g16535, g17590, g17616, g17700, g17949, g22276, g22280, g22289, g22304, g22306, g22308, g22310, g22315, g22331, g22332, g22338, g22346, g22348, g22357, g22367, g22372, g22476, g22485, g22497, g25077, g25094, g31340, g6647, g6926 |
| GO:0055085 | transmembrane transport | 4.4E-02 | 4.4E-02 | g14478, g22276, g22280, g22306, g22308, g22315, g22331, g22338, g22346, g22485, g25077 |

### Variation in gene age along chromosomes reveals that evolutionarily young loci produce abundant small RNAs

To investigate genome-wide patterns of feature distribution, a series of datasets were mapped to *R. irregularis* chromosomal scaffolds using non-parametric linear regressions. Normalised reads per kilobase per million reads mapped (RPKM) of Nanopore RNA-Seq (full-length, poly(A)-selected) and small RNA-Seq (~24nt long) were reported in 200bp genomic bins, and gene ages (pre-HDF test) were measured across chromosomal length (**Figure 5**). An uneven distribution of gene age was observed for all chromosomes, distinguishing regions enriched with evolutionarily ancient genes (low phyloranks) from regions with evolutionarily young genes (high phyloranks) (**Figure 5**). Genomic regions with evolutionarily ancient genes tend to have high poly(A)+ RNA and low small RNA expression levels, and these patterns are reversed in regions with evolutionarily young genes (**Figure 5** and **S3ABD**). However, a small number of genes produce small RNAs (Dallaire et al., 2021), and when examined at the scale of individual genes, per-gene small RNA expression levels did not correlate with gene age (**Figure S3C**). This led to the conclusion that sequence surrounding young genes, rather than young genes themselves, drive the observed pattern of small RNA expression. Two particular loci of ~2Mbps in length were identified (**Figure 5**, regions shaded in blue) that collectively contain evolutionarily young coding regions and the most abundant concentration of highly expressed small RNA loci. These data suggest that the genome of *R. irregularis* presents highly transcribed regions harbouring highly conserved genes, and lesser transcribed, small RNA-producing regions with evolutionarily younger genes.

**Figure 5.**
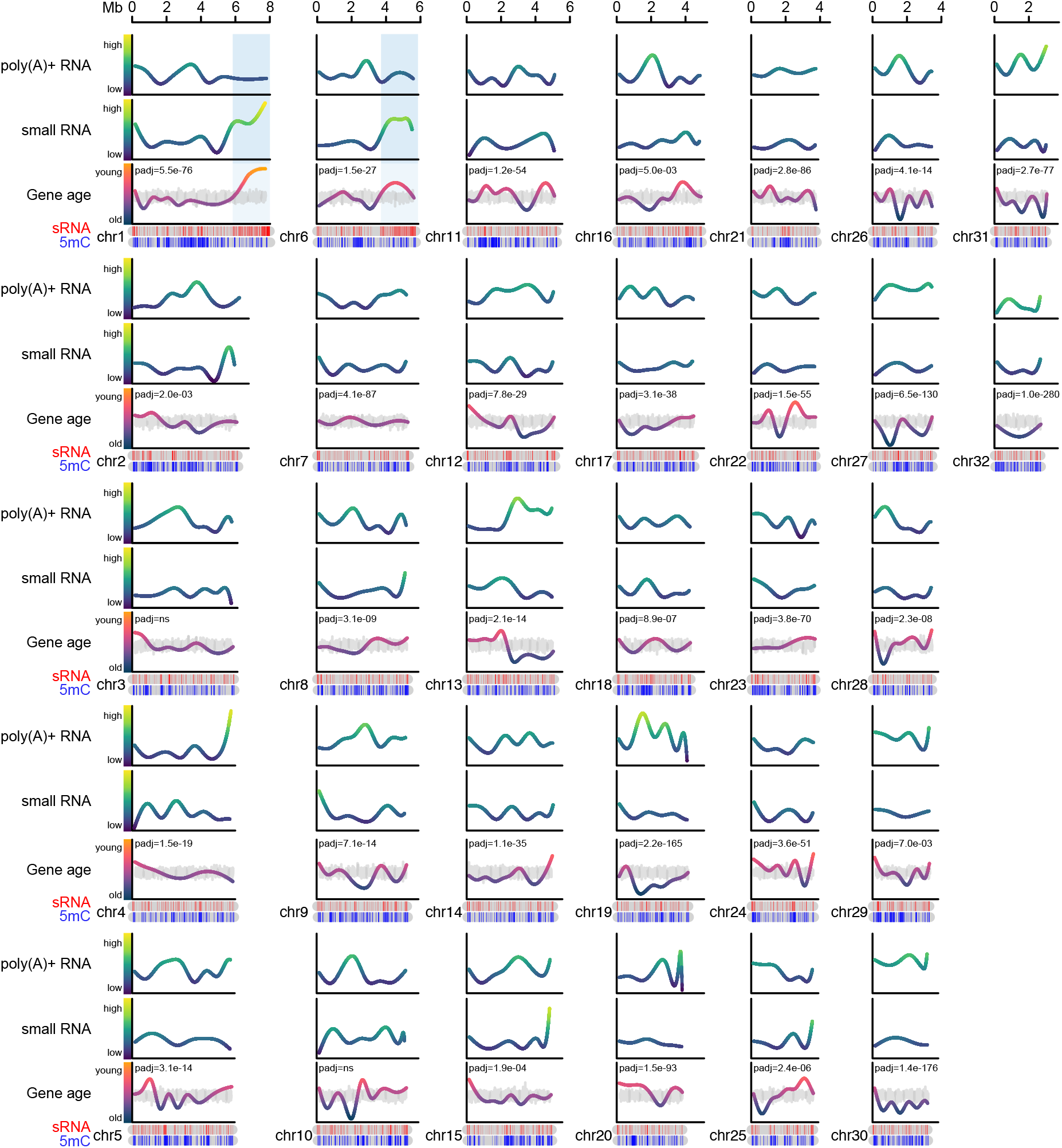
Genome-wide patterns of gene age and expression. Per-chromosome genomic distribution of Nanopore polyA+ RNA-Seq (top line graph; long RNA), expression of small non-coding RNAs (RPKM, second line graph; sRNA), gene age (third line graph; old to young corresponding to phyloranks 1 to 9), small RNA loci (top ideogram; sRNA) and highly methylated CGs (bottom ideogram; 5mC values > 80% are shown; 5mC). Colour gradients on line graphs match the y-axis scales. Grey line graphs overlapping with gene ages represent the mean of 1000 random permutations of gene ages, and a paired t-test (grouped by chromosome) was used to test the significance of observed gene age distributions relative to random permutations (padj). Values used for non-parametric linear regressions of long and small RNA expression are normalised RPKM calculated in 200bp bins. Gene ages are regressed and plotted following chromosomal gene distribution (not binned).

### Discussion

The number of fungal species with highly contiguous, long-read, and chromosome-scale assemblies lags behind that of animals and plants (Marks et al., 2021; Rhie et al., 2021). This work presents a chromosome-scale assembly of the symbiotic fungus *R. irregularis*, isolate DAOM197198, the model species for molecular research into AM fungi. This assembly of 32 chromosomal scaffolds is highly contiguous, with only 10 gaps and a contig N50 of 3.9Mb. Nuclear chromosomes display a very high synteny with those of a previous assembly of *R. irregularis* DAOM197198 (**Figure S1C**) (Yildirir et al., 2022), though this assembly assigns sequence to 32 chromosomal scaffolds, in contrast to the 33 chromosomal scaffolds previously presented. A complete, gapless, circular mitochondrial genome of 70,793 bp was also assembled, a size consistent with a previous assembly of the *R. irregularis* mitochondria (Lee & Young, 2009). This novel assembly, alongside a high-quality genome annotation of *R. irregularis*, consisting of gene models with corrected structures, splice junctions and untranslated regions, will further aid research into *R. irregularis* and AM fungal biology, as well as comparative genomics approaches.

This highly contiguous genome assembly enabled an analysis of chromosomal distributions of *R. irregularis* genomic features and gene and small RNA expression. This supports a previous observation of functional and evolutionary genome compartmentalisation in *R. irregularis* (Yildirir et al., 2022), and build on this work by showing that chromosomes contain highly expressed regions with highly conserved genes, and lowly-expressed regions hosting more recently evolved genes. This is reminiscent of the two-speed genome model, which has been described in filamentous phytopathogens (Torres et al., 2020) and proposed to exist in AM fungi (Reinhardt et al., 2021; Yildirir et al., 2022). According to this model, fast-evolving virulence-associated genes are compartmentalised into repeat-rich genomic regions or accessory chromosomes that are depleted of conserved housekeeping genes. In the plant pathogenic fungus *Sclerotinia sclerotiorum*, small RNAs originate from transposable elements in polymorphic genome compartments (Derbyshire et al., 2019). In *R. irregularis*, quantitative evidence for differential evolutionary speed and sequence variation in genomics compartments is lacking. Nevertheless, evolutionary patterns of genomic architecture can be observed, as well as small RNA production in regions with evolutionarily young coding spaces (**Figure 5**). Genome-wide patterns of small RNA expression may therefore point to loci that encode the basis for lineage-specific adaptations and diversification in AM fungi, and may facilitate studies into adaptive structural and sequence variation.

With the increasing number of reference genomes available for Earth’s biodiversity (Lewin et al., 2018) and the development of efficient algorithms for sequence analysis (Buchfink et al., 2021; Jumper et al., 2021), characterisation of genes and genomes can harness comparisons at tree-of-life scale. This study used a phylostratigraphic gene age inference tool that performs alignments against the entire NCBI non-redundant protein database to trace back the emergence of *R. irregularis* genes (Barrera-Redondo et al., 2022). Genetic machinery for phosphate, ammonium, monosaccharide transport, ion transmembrane transport and a group of transmembrane ion channels were found to have evolved at or before the Mucoromycota phylorank, thereby predating the emergence of Glomeromycotina. Genes encoding key symbiotic functions of AM fungi therefore already existed in the Mucoromycota ancestor. A similar phenomenon was observed in the genomes of saprotrophic fungi, which encode the symbiosis toolkit of their successor ectomycorrhizal species (Hess et al., 2018; Miyauchi et al., 2020). Similarly in plants, the genetic basis for symbiont perception, nodule organogenesis and nitrogen-fixation genes already existed in the common ancestor of nitrogen-fixing legumes, and diversified in downstream nitrogen-fixing lineages (Libourel et al., 2022). Such macroevolutionary transitions punctuate the eukaryotic tree of life, where the acquisition of new molecular functions accompanies major evolutionary and ecological transitions, but precedes divergence and lifestyle specialisation in downstream lineages (Domazet-Lošo et al., 2022; Ocana-Pallares et al., 2022).

The detection of a gene birth event accompanying the emergence of Glomeromycotina highlights the existence of previously undescribed lineage-restricted innovation. Gene birth events associate with the emergence of ectomycorrhizal lifestyles (Hess et al., 2018; Miyauchi et al., 2020) and of rhizoid and root development in land plants (Barrera-Redondo et al., 2022). While the loss of *de novo* fatty acid synthase activity likely played a major role in creating dependence to externally supplied carbon (Keymer et al., 2017; Luginbuehl et al., 2017; Malar et al., 2021; Trepanier et al., 2005), the birth of lineage-specific genes such as the transcription factors identified here may also underlie an evolutionary transition in the Glomeromycotina subphylum.

## Materials and methods

### DNA preparation and sequencing

High molecular weight (HMW) DNA was extracted from 2g of *R. irregularis* DAOM197198 Grade A spores (Agronutrition) (Schwessinger & McDonald, 2017). 100mg of ground spore material was resuspended in lysis buffer and processed as indicated. Two successive rounds of cleanup were performed using a 0.45X volume of Ampure XP beads in DNA-Lo-Bind tubes following the Manufacturer’s protocol. DNA was finally eluted in 50uL of 10mM Tris-pH8. DNA quality was assessed by running on a 0.5% agarose gel. Sequencing libraries were prepared using the Oxford Nanopore Rapid DNA sequencing kit SQK-RAD004 and sequenced on MinION flow cells FLO-MIN106D following the accompanying protocol. Genomic Nanopore reads were basecalled with Guppy Basecalling Software version 5.0.11+2b6dbff (Oxford Nanopore Technologies, Limited).

### *R. irregularis* DAOM197198 genome assembly and polishing

9.06 Gb of Nanopore sequence reads were trimmed to remove adapters using Porechop (version 0.2.4), and 1,288,465 of 1,288,893 reads were retained after trimming (99.97%). Following trimming, read N50 was 24,957 bp. The Shasta long-read assembler (shasta-Linux-0.8.0) was then used to produce a raw genome assembly using the parameters ­­Assembly.consensusCaller Bayesian:guppy-5.0.7-a, --Kmers.k 10, --MinHash.minHashIterationCount 50, --Align.bandExtend 20, -- Align.downsamplingFactor 0.1, --ReadGraph.creationMethod 0, --ReadGraph.maxAlignmentCount 12, --ReadGraph.crossStrandMaxDistance 0, --Align.minAlignedFraction 0.3, -- Align.minAlignedMarkerCount 60, --Align.maxSkip 50, --Align.maxDrift 30, --Align.maxTrim 30, -- MarkerGraph.minCoveragePerStrand 3, --Assembly.iterative, and --Assembly.pruneLength 1500.

The raw assembly was then trimmed of contigs smaller than 500bp (removing two contigs). Subsequent polishing of this trimmed assembly was carried out using the PEPPER-Margin-DeepVariant pipeline as described in (Shafin et al., 2021; Shafin et al., 2020). Broadly, the Nanopore reads described above were aligned against the raw, trimmed *R. irregularis* assembly using minimap2 (parameters: -ax map-ont). 83.7 Gb of Illumina reads obtained from (Maeda et al., 2018) were also aligned against this assembly using BWA-MEM with default parameters (Li, 2018). Alignments of the Nanopore and Illumina reads produced variant calls that were corrected in the assembly using the PEPPER-Margin-DeepVariant pipeline and Merfin (Formenti et al., 2022).

To assemble the telomeric regions of this genome, 1964 reads containing the telomeric repeat TTAGGG_8_ were extracted from trimmed Nanopore reads. These repeat-containing reads were then used to assemble 62 telomeric-contigs using Shasta with parameters as described above, with the exception of --Assembly.consensusCaller Bayesian:guppy-5.0.7-a and --Kmers.k 14. The 62 telomeric contigs were polished using the same polishing pipeline as described above, mapping the initial telomere repeat-containing reads and genomic Illumina reads to the telomeric contigs and polishing using the PEPPER-Margin-DeepVariant pipeline. The full genome contigs and the telomeric contigs were then manually fused based on overlapping sequence identified following minimap2 alignment (parameters: -ax map-ont). The QV score of the raw assembly was Q29.49, rising to Q32.6 following polishing with PEPPER, and finally Q36.27 after polishing with DeepVariant and fusing with separately assembled and polished telomeric contigs.

The assembly process resulted in the assembly of a complete, circular mitochondrial genome of 70,793bp. Circularity of the mitochondrial assembly graph was visualised using Bandage (**Figure S1A**) (Wick et al., 2015). MitoHifi v.2.2 (Allio et al., 2020; Laslett & Canback, 2008; Uliano-Silva et al., 2021) was used to annotate the mitochondrial genome (**Figure 1A**). This mitochondrial genome was removed from the nuclear genome assembly for manual curation. Hi-C read data for *R. irregularis* DAOM197198 (Yildirir et al., 2022) was aligned to the remaining 42 contigs using BWA-mem (Li & Durbin, 2009) and the subsequent alignment file was used to produce a PretextView Map (Harry, 2020). The PretextView Hi-C contact map was manually curated and the assembled contigs were pieced together to produce chromosome-scale scaffolds (Howe et al., 2021).

### Quality assessment of the *R. irregularis* DAOM 197198 assembly

The genome assembly was scored by BUSCO version 5.2.2 (Simao et al., 2015) as 95.8% complete using the fungi_odb10 database. 726 complete BUSCOs were identified out of a total of 758 BUSCO groups searched, of which 13 were duplicated. All trimmed Nanopore reads were mapped to the assembly using Minimap2 (parameters: -ax -map-ont) (Li, 2018), resulting in the mapping of 1,279,771 reads to the final assembly (99.32% of total trimmed reads). Mosdepth was used to examine the cumulative distribution of read coverage for each contig. Average Nanopore read coverage was highly uniform; between 77 and 85X across all nuclear contigs (**Figure S1B**). To assess Illumina read coverage uniformity, BWA-MEM was used to align Illumina genomic DNA reads (Maeda et al., 2018) to the assembly, with 200,768,646 of 211,520,841 (94.61%) reads mapping successfully. Average Illumina read coverage of contigs identified through mosdepth was again uniform, all contigs displayed coverage between 227X and 232X (**Figure S1B**). Additionally, a BLASTn analysis (parameters: -task megablast, -max_target_seqs 25, -culling_limit 2, -evalue 1e-25) was carried out on the assembly and the best hit for each contig was *R. irregularis.*

### Genome annotation

*R. irregularis* DAOM197198 RNA samples (plates of 50,000 spores/sample) used for genome annotation and the protocol for the production of rice exudates to treat spore plates were the same as described previously (Dallaire et al., 2021). The Illumina RNA-Seq samples used were an untreated spore plate, a 24h rice exudate-treated sample, a 48h rice exudate-treated sample (48e_1), and a sample of *R. irregularis* colonised maize root (growth conditions with RNA extraction as described for rice plants in (Dallaire et al., 2021). Additional Illumina RNA-Seq samples from a further experiment described in the same publication used for genome annotation were two *Nicotiana benthamiana* root samples colonised by *R. irregularis*, and two germinated spore samples. Short-read library preparation, sequencing and adapter trimming were carried out on paired-end polyA+ RNA by Novogene UK Co. Ltd. with read lengths of 150bp.

The TrimGalore!-0.6.6 wrapper script for Cutadapt (Martin, 2011) was used for quality and adapter trimming of all short-read fastq files (parameters: --length 36 -q 20 --stringency 1 -e 0.1 --paired -- phred33).). For alignment of the Illumina RNA-Seq files to the soft-masked genome assembly, STAR (version 2.7.6a) was used (parameters: --outFilterMultimapNmax 20) (alignment statistics in Table 2) (Dobin et al., 2013). Output BAM files from this STAR alignment were used as input for BRAKER 2.1.5 (parameters: –gff3 –fungus –softmasking) (Bruna et al., 2021). Protein domains were predicted from BRAKER2 models using InterProScan 5.55-88.0 (Jones et al., 2014), and were manually curated to remove genes with transposon-related protein domains, leading to the Illumina-based gene annotation presented in this study.

To produce the long-read RNA-Seq data used to refine Illumina-based gene models, Nanopore RNA-Seq was carried out using a sample of *R. irregularis* DAOM197198 pre-germinated spore plate (50,000 spores/plate) (Table 2 and described below). 1ug of total RNA from three samples were individually poly(A) selected using the NEBNext Poly(A) mRNA Magnetic Isolation Module (NEB #E7490). Poly(A)+ concentration and rRNA depletion were assessed using Qubit and TapeStation. ~20ng from each sample (total of 70ng of poly(A)-selected RNA) was barcoded using the Nanopore PCR-cDNA barcoding kit (Kit SQK-PCB109). PCR was performed with 14 cycles and 6.5 minute extensions. 33.3 fmol of each barcoded sample were pooled together and prepared for sequencing on an R9.4.1 flow cell. Sequence reads were demultiplexed and basecalled using Guppy Basecalling Software version 5.0.11+2b6dbff (Oxford Nanopore Technologies, Limited). Following basecalling, Pychopper version 2.5 was used to trim the reads and rescue fused reads. These reads were provided as evidence to the update script of funannotate version 1.8.7 (https://zenodo.org/record/4054262#.Yv4hJy8w3AY), which uses PASA (Haas et al., 2003; Haas et al., 2008), to refine gene models of the Illumina-based gene annotation, to predict 5’UTR, 3’UTR and polyadenylation signal sequences, and to extract poly(A) tail sequences. Transposon-related protein domains were removed from the updated gene models, leading to the final Illumina+Nanopore-based gene annotation presented in this study.

Illumina-based and Illumina+Nanopore-based gene models were functionally annotated separately using funannotate version 1.8.7, ran with the BUSCO database “fungi” and the UniProt DB version 2022_01, and with protein domain prediction evidence from 1) InterProScan 5.55-88.0 (Jones et al., 2014), 2) eggnog-mapper v2.1.7 (with more-sensitive mode, corresponding to diamond.2.0.8’s very-sensitive mode (Buchfink et al., 2021; Cantalapiedra et al., 2021; Huerta-Cepas et al., 2019), and 3) Secondary metabolism and transmembrane domain prediction using antiSMASH version 6.0.1 (Blin et al., 2021). Gene annotations were scored by BUSCO with the fungi_odb9 database. GO terms associated with the Illumina+Nanopore-based annotation were processed with g:Profiler’s GMT tool, and GO term analyses were performed using the g:Profiler web server (Raudvere et al., 2019) using the token “gp__xfGY_dQeI_yx4”, or the GMT file provided as Supplemental File.

### Repeat and transposable element annotation

Repeats were modelled using EDTA (parameter --sensitive 1)(Ou et al., 2019). Protein domains were predicted from consensus sequences using InterProScan 5.55-88.0. Consensus sequences containing gene-related protein domains were filtered out, and the remaining sequences were used to annotate the genome using RepeatMasker (parameters -s -no_is -norna -nolow -div 40) (Smit et al., 2015).

### Small RNA annotation and quantification

70,956,710 small RNA-Seq reads from two replicates of oxidised and two replicates of column-purified spore RNA (Dallaire et al., 2021) were used to run ShortStack (Axtell, 2013) (parameters --dicermin 20 --dicermax 27 --foldsize 300 --pad 200 --mincov 10.0rpm --strand_cutoff 0.8 --mmap r).

### DNA methylation basecalling

161Gb of raw FAST5 files obtained from three R9.4.1 flow cells were basecalled with Guppy5, producing 985,449 reads which were successfully processed by tombo (Stoiber et al., 2017) and used by DeepSignal2 (Ni et al., 2019) to extract CG motifs and to call modifications using a human model (model.dp2.CG.R9.4_1 D.human_hx1.bn17_sn16.both_bilstm.b17_s16_epoch4.ckpt.

### Phylostratigraphy analyses

GenEra (Barrera-Redondo et al., 2022) was run using DIAMOND in ultra-sensitive mode (Buchfink et al., 2021). An E-value threshold of 1E^-5^ was chosen to balance the detection of distant homologs while minimizing the amount of false positives against the NR database (Barrera-Redondo et al., 2022). Taxonomy IDs used for the focal species are 50956 for *G. pyriformis*, 4874 for *G. margarita*, 1432141 for *R. irregularis*, 101101 for *D. decumbens*, 1314771 for *M. elongata*, 64574 for *R. spectabilis* and 4837 for *P. blakesleeanus*. Genes with taxonomic representativeness scores below 30% were flagged as possible contamination or horizontal gene transfer and were not included in subsequent analyses. Some phyloranks were corrected: strain level ranks were moved to species level (*“R. irregularis* DAOM 197198*”* to *“R. irregularis” and “Linnemannia elongata AG-77” to “Linnemannia elongata”)* and *“*Fungi incertae sedis*”* was moved to the kingdom level “Fungi”. Several phyloranks were collapsed due to unresolved phylogenetic placement of subphyla (Figure S2B-D) or insufficient genomic data (Figure S2A). GenEra’s homology detection failure test (Barrera-Redondo et al., 2022; Weisman et al., 2020) was run by using the pairwise evolutionary distances from a phylogenomic tree (Li et al., 2021) to obtain a list of genes in *R. irregularis* whose ages cannot be explained by gene untraceability from the genus to the kingdom phyloranks.

### Chromosomal distribution of genomic features and expression

Nanopore RNA-Seq reads were trimmed of adapters and cleaned with seqclean (Chen et al., 2007) to remove a percentage of undetermined bases, polyA tails, overall low complexity sequences and short terminal matches. Cleaned sequences were then mapped using minimap2 (options -G max intron length=3000, -ax, map-ont)(Li, 2018). Small RNA-Seq reads were aligned to the genome using bowtie (options --mmap r) (Langmead & Salzberg, 2012). Nanopore and small RNA RPKM were calculated using bamCoverage (options –bam --binSize 200 --ignoreDuplicates --normalizeUsing RPKM) (Ramirez et al., 2016). A general additive model was used to regress feature values across chromosome lengths (gam(<gene age or RPKM> ~ s(Chrom.start, bs = “cs”, by=Chrom)), using the package mgcv v.1.8-40. Gene age fits were plotted using the start position of each gene and RPKM fits were plotted using the start position of every 200bp bin. Gene ages were randomly permuted 1000 times and the mean was plotted using the unchanged start position of each gene. A paired t-test grouped by chromosome was used to test the significance of the observed gene age distributions relative to random permutations.

## Data availability

DNA and RNA sequencing datasets generated are available at PRJNA885267. Previously published datasets used are GSE172187 and PRJNA722386 from (Dallaire et al., 2021), PRJNA748024 from (Yildirir et al., 2022) and DRA004835 from (Maeda et al., 2018). Code and Supplemental Files are available at https://github.com/bethanmanley.

## Author contributions

B. F. M.: Conceptualization, Formal analysis, Investigation, Visualization, Methodology, Writing — original draft, Writing — review and editing.

J. S. L.: Formal analysis, Investigation, Writing — review and editing.

J. B-R.: Formal analysis, Investigation, Writing — review and editing.

G.Y.: Resources, Writing — review and editing.

J. S.: Writing — review and editing.

N. C.: Writing—review and editing. U. P.: Writing — review and editing.

E. A. M.: Funding acquisition, Writing — review and editing.

A. D.: Conceptualization, Formal analysis, Investigation, Visualization, Methodology, Project administration, Writing—original draft, Writing—review and editing.

## Acknowledgements

We thank Hajk-Georg Drost, Susana Coelho and members of their groups, as well as all members of the Miska and Paszkowski laboratories for providing valuable input on the manuscript. We thank Jen McGaley for producing and sharing the image shown in Figure 1. We thank Uku Raudvere and Hedi Peterson for helping with g:Profiler, and David Jordan for helping with data analysis and statistics. We thank Paolo Carnevali and Kishwar Shafin for helping with genome assembly and polishing, and Charles Bradshaw for bioinformatics support. We thank the Tree of Life consortium for sharing genome curation pipelines.

## Supplemental Information

Supplemental Data and Files are available at BioProject PRJNA885267 and https://github.com/bethanmanley.

## Supplemental Files

### Raw Nanopore DNA and RNA reads (BioProject PRJNA885267)

Rhizophagus_irregularis_DAOM197198_Nanopore_DNASeq_Run1.fastq.gz
Rhizophagus_irregularis_DAOM197198_Nanopore_DNASeq_Run2.fastq.gz
Rhizophagus_irregularis_DAOM197198_Nanopore_DNASeq_Run3.fastq.gz
Rhizophagus_irregularis_DAOM197198_Nanopore_RNASeq.fastq.gz

### Nuclear genome assembly, masked and unmasked

Rhizophagus_irregularis_DAOM197198_assembly.fasta
Rhizophagus_irregularis_DAOM197198_assembly_masked.fasta

### Illumina and Illumina+Nanopore genes, CDS, mRNA, protein, and functional annotations

Rhizophagus_irregularis_DAOM197198_Illumina+ONT_curated.gff3
Rhizophagus_irregularis_DAOM197198_Illumina_curated.gff3
Rhizophagus_irregularis_DAOM197198_annotations_Illumina+ONT.txt
Rhizophagus_irregularis_DAOM197198_annotations_Illumina.txt
Rhizophagus_irregularis_DAOM197198_cds-transcripts_Illumina+ONT_curated.fa
Rhizophagus_irregularis_DAOM197198_cds-transcripts_Illumina_curated.fa
Rhizophagus_irregularis_DAOM197198_mrna-transcripts_Illumina+ONT_curated.fa
Rhizophagus_irregularis_DAOM197198_mrna-transcripts_Illumina_curated.fa
Rhizophagus_irregularis_DAOM197198_proteins_Illumina+ONT_curated.fa
Rhizophagus_irregularis_DAOM197198_proteins_Illumina_curated.fa

### GO terms for g:Profiler

Rhizophagus_irregularis_DAOM197198_Illumina+ONT_GOterms.gmt

### Repetitive and transposable element annotation

Rhizophagus_irregularis_DAOM197198_curatedrepeatlibrary.fasta
Rhizophagus_irregularis_DAOM197198_repeatmasker.out
Rhizophagus_irregularis_DAOM197198_repeats.gff3

### DNA methylome (spores)

Rhizophagus_irregularis_DAOM197198_mCG_mods_frequency.tsv

### Poly(A) signal and tail sequences

Rhizophagus_irregularis_DAOM197198_pasa_polyAsite_analysis.out
Rhizophagus_irregularis_DAOM197198_pasa_polyAsites.fasta

### Small RNA annotation

Rhizophagus_irregularis_DAOM197198_sRNA.gff3
Rhizophagus_irregularis_DAOM197198_sRNA.tsv

### Mitochondrial genome assembly and annotation

Rhizophagus_irregularis_DAOM197198_mtDNA.fasta
Rhizophagus_irregularis_DAOM197198_mtDNA.gff

### *R. irregularis* phylostratigraphy

Rhizophagus_irregularis_DAOM197198_1432141_phyloranks.tsv
Rhizophagus_irregularis_DAOM197198_1432141_high-confidence_phyloranks.tsv

### Mucoromycota fungi phylostratigraphy

Disdec1 _101101_phyloranks.tsv
Geopyr1_50956_phyloranks.tsv
Gigmar1_4874_phyloranks.tsv
Morel2_1314771_phyloranks.tsv
Phybl2_4837_phyloranks.tsv
Radspe1_64574_phyloranks.tsv

---

**Supplemental Figure 1.**
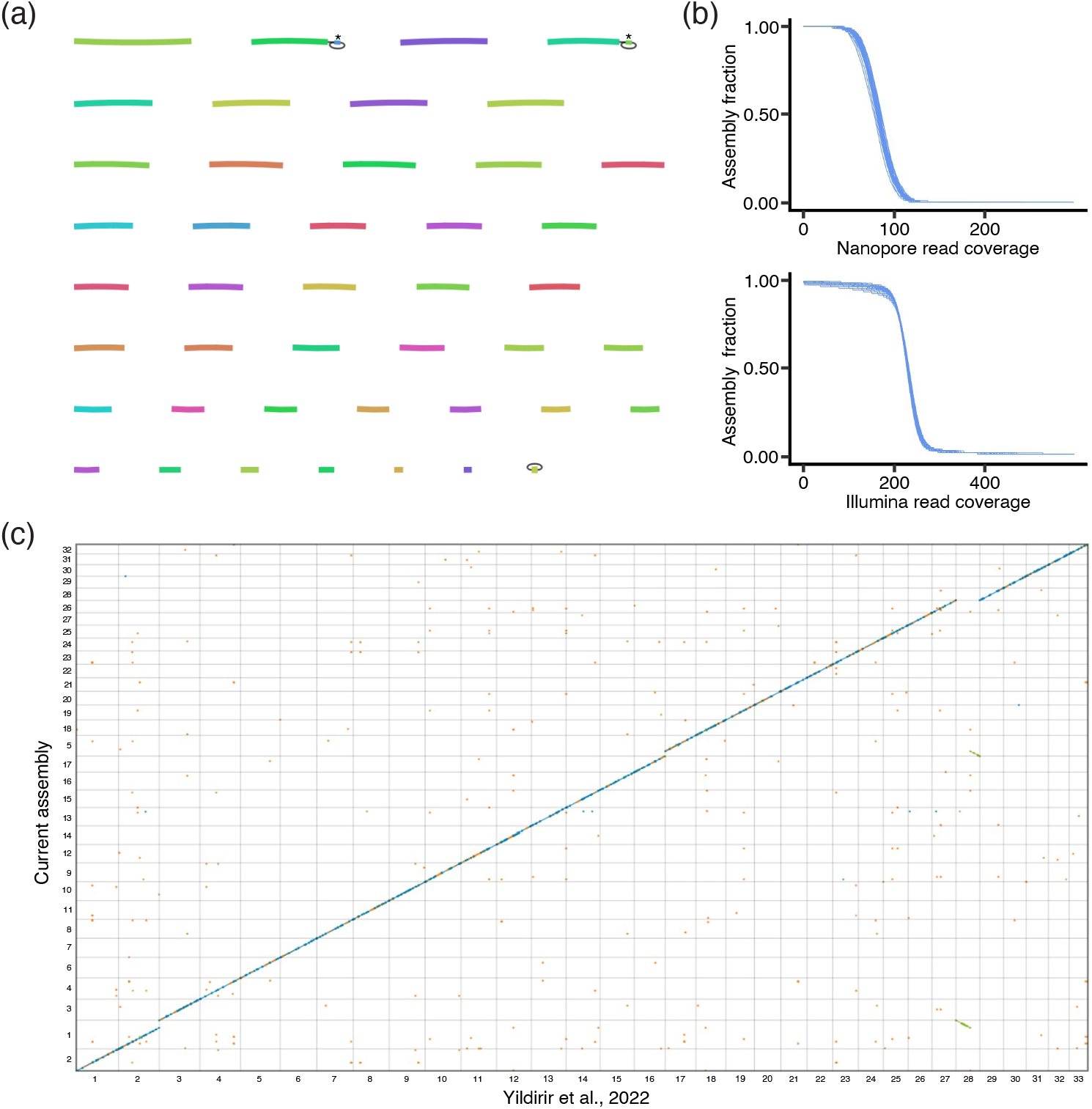
Assembly graph, coverage and macro-synteny analysis. a) Raw assembly graph following Shasta assembly. Ambiguities in the graph structure are represented by edges (fine black lines). Contigs indicated with a star (*) have lengths of 212bp and 164bp and were removed from the assembly due to their size. The circular mitochondrial contig is displayed as the final contig. b) Cumulative Nanopore (top) and Illumina (bottom) read coverage for each of the assembled contigs, pre-curation. c) Whole-genome pairwise alignment between the current assembly and the Yildirir et al., 2022 assembly. Blue dots represent unique alignments on the positive strand of both assemblies, green dots represent unique alignments on the negative strand, and orange dots represent non-unique alignments (repetitive).

**Supplemental Figure 2.**
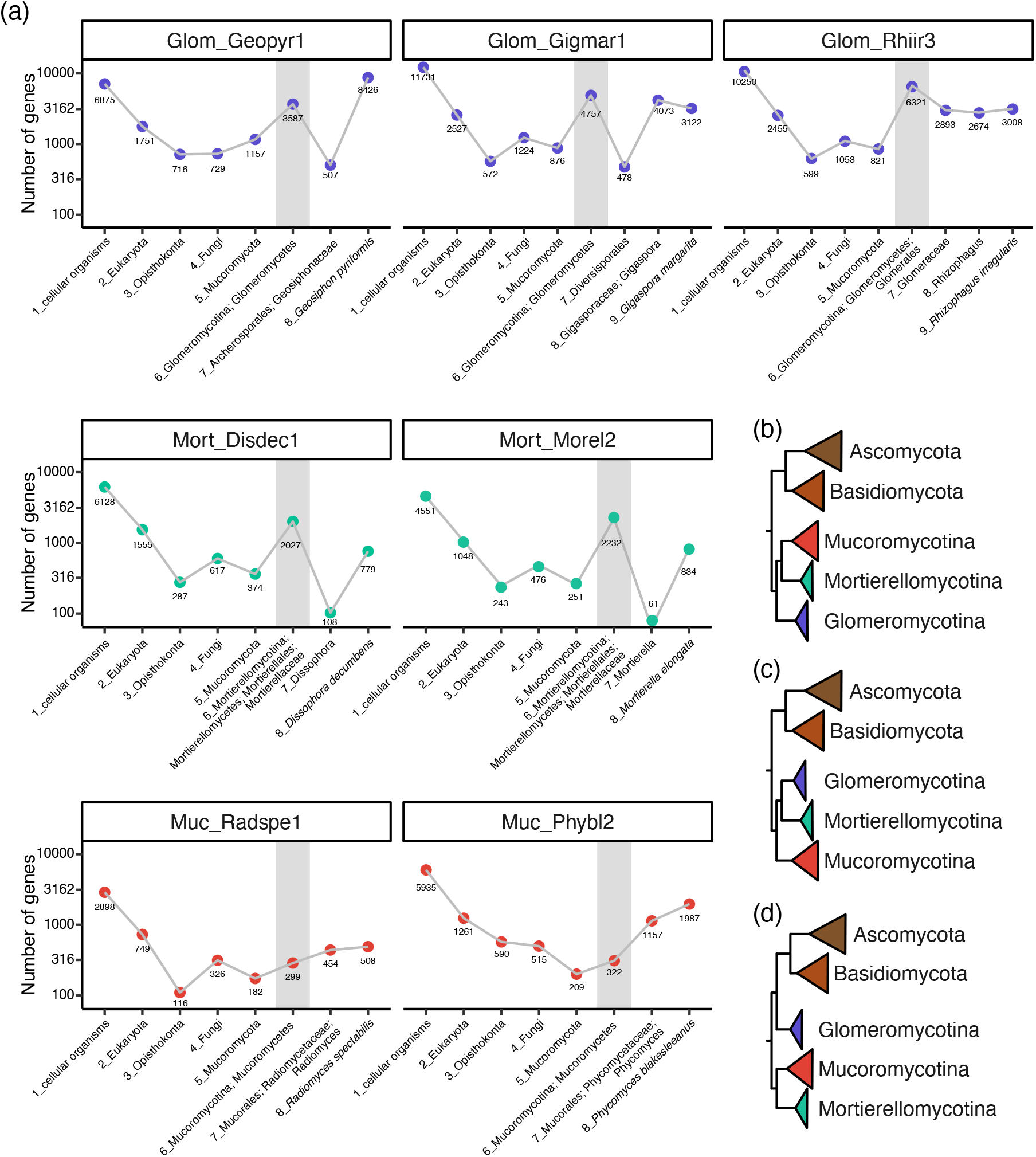
Distribution of gene age in Mucoromycota fungi. a) Gene age assignment for Glomeromycotina (*Geosiphon pyriformis, Gigaspora margarita* and *Rhizophagus irregularis)*, Mortierellomycotina (*Dissophora decumbens* and *Mortierella elongata)* and Mucoromycotina (*Radiomyces spectabilis* and *Phycomyces blakesleeanus)* species. Log-transformed number of genes in each phylorank is displayed. Grey shaded areas highlight phyloranks that were collapsed due to insufficient genomic data, which likely result in overestimation of gene birth at phylorank 6 for the Glomeromycotina and Mortierellomycotina subphyla. b), c) and d) show possible phylogenetic placement of the three subphyla.

**Supplemental Figure 3.**
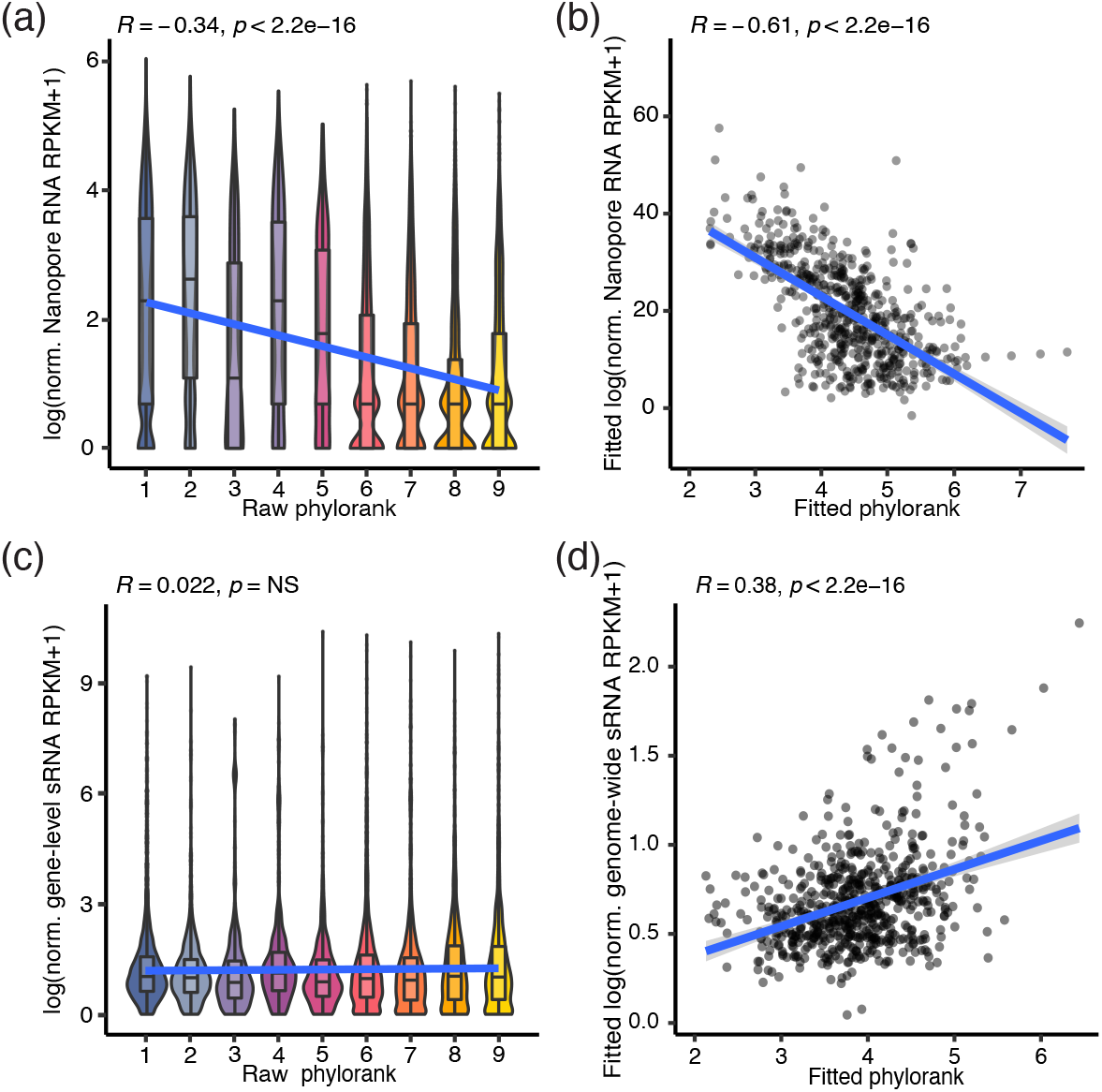
Correlation of genomic feature distributions. a) Distribution of raw values of normalised Nanopore RNA RPKM, in 200bp bins, across phyloranks. b) Scatter plot showing linear regression values of normalised Nanopore RNA RPKM and phylorank values, for a subsample of 200bp genomic bins. c) Distribution of raw values of normalised small RNA RPKM measured at gene level, across phyloranks. d) Scatter plot showing linear regression values of normalised genome-wide small RNA RPKM and phylorank values, for a subsample of 200bp genomic bins.

